# Single cell RNA Sequencing Identifies G-protein Coupled Receptor 87 as a Novel Basal Cell Marker of Distal Honeycomb Cysts in Idiopathic Pulmonary Fibrosis

**DOI:** 10.1101/2021.09.03.458324

**Authors:** Katharina Heinzelmann, Qianjiang Hu, Yan Hu, Evgenia Dobrinskikh, Henrik M. Ulke, Meshal Ansari, Colton Leavitt, Carol Mirita, Tammy Trudeau, Maxwell L. Saal, Pamela Rice, Bifeng Gao, William J. Janssen, Ivana V. Yang, Herbert B. Schiller, Eszter K. Vladar, Mareike Lehmann, Melanie Königshoff

## Abstract

Idiopathic Pulmonary Fibrosis (IPF) is a progressive and fatal lung disease with limited therapeutic options. Epithelial reprogramming and honeycomb cysts are key pathological features of IPF, however, the IPF distal bronchiole cell subtypes and their potential contribution to IPF development and progression still remain poorly characterized. Here, we utilized single-cell RNA sequencing on enriched EpCAM^+^ cells of the distal IPF and Donor lung. Using the 10x Genomics platform, we generated a dataset of 47,881 cells and found distinct cell clusters, including rare cell types, such as suprabasal cells recently reported in the healthy lung. We identified G-protein coupled receptor (GPR) 87 as a novel surface marker of distal Keratin (KRT)5^+^ basal cells. GPR87 expression was localized to distal bronchioles and honeycomb cysts in IPF in situ by RNA Scope and immunolabeling. Modulation of GPR87 in primary human bronchial epithelial cells cultures resulted in impaired airway differentiation and ciliogenesis. Thus, GPR87 is a novel marker and potentially druggable target of KRT5^+^ basal progenitor cells likely contributing to bronchiole remodeling and honeycomb cyst development in IPF.

## Introduction

Idiopathic Pulmonary Fibrosis (IPF) is a devastating and life-threatening lung disease characterized by epithelial reprogramming and increased extracellular matrix deposition leading to loss of lung function. A prominent histopathological structure in the IPF lung are aberrant bronchioles filled with mucus, referred to as honeycomb cysts[1]. These heterogeneous bronchiolized areas feature clusters of simple epithelium with Keratin (KRT)5^+^ basal-like cells interspersed with pseudostratified epithelium containing differentiated, hyperplastic epithelial cells as well as aberrant ciliated cells[2-5]. Recent single-cell RNA sequencing studies of the lung epithelium shed further light into cellular subtypes unique to IPF, including basaloid KRT5^-^/KRT17^+^ cells present in the distal lung[6-9]. However, IPF distal bronchiole cell subtypes still remain poorly characterized and their disease contribution remains underinvestigated. Using single-cell RNA sequencing of enriched EpCAM^+^ lung cells of the distal IPF lung, we identified *G-protein coupled receptor (GPR) 87*, as a novel marker of distal bronchioles and KRT5^+^ IPF basal cells contributing to impaired airway differentiation and honeycombing.

## Material and Methods

### Human lung material

Fresh non-fixed human lung tissue from de-identified healthy donors and IPF explanted patient donors was received from National Jewish Hospital/UC Health University of Colorado Hospital (Denver, CO, USA) (COMIRB 11-1664).

### Single-cell RNA sequencing

Human lung tissue (IPF/age-matched donors, each n=3) was homogenized and 4 g of tissue were digested by dispase/collagenase (Collagenase: 0.1U/mL, Dispase: 0.8U/mL, Roche) for 1 hour at 37°C. Samples were successively filtered through nylon filters (100 µm and 20 µm) followed by a percoll gradient and MACS sorting with CD45 Microbeads (Miltenyi Biotec). Cells were stained with APC anti-human EpCAM antibody (Biolegend) and DAPI (ThermoFisher Scientific) and subjected to FACS. Single-cell suspensions were loaded onto a Chromium single cell chip (Chromium™ Single Cell 3’ Reagent Kit, v2 Chemistry) to obtain single cell 3’ libraries for sequencing. The barcoded libraries were sequenced using Illumina HiSeq 4000 (Illumina, San Diego, CA, USA). Raw reads were aligned to the hg19 reference genome. The count matrix of each sample was integrated and pre-processed using Scanpy (v1.4.6). In details, BBKNN (v1.4.0) was used to regress out the batch affection. Cells with less than either 1000 counts or 400 genes and cells with more than 15% mitochondria reads were filtered out. Data analysis was performed using Python and R packages, including Scanpy (v1.4.6), EnhancedVolcano (v1.10.0), pheatmap (v1.0.12) and ggplot2 (v 3.3.3).

### Cell culture and qPCR

Primary human bronchial epithelial cells (HBECs) were isolated from healthy donors (n=3) as described[4] and seeded on 6 cm dishes within BEGM media (Lonza) before transduction with lentivirus (empty vector (Origene, PS100092) or human GPR87 ORF (Origene, RC218486L3)) for 6 hrs. Medium change was followed by 24 hrs puromycin selection. Cells were transferred and kept submerged on transwells till reaching an intact tissue structure. Cells were set on air liquid interface (ALI) (=day 0) using PneumaCult ALI media (STEMCELL Technologies) and cultured until day 21 or 28 with media change every other day. ALIs were stimulated every other day with TGF-β1) (R&D, 240-B-002, 2 or 4 ng/ml) starting day 21. RNA was isolated using the RNeasy Plus Kit (Qiagen). cDNA was synthesized and qPCR performed using SYBR Green PCR master mix at a Roche LightCycler 96. Primers used: huGPR87-fw (ACCTATGCTGAACCCACGC), -re (CCGTGCAGCTCGTTATTTGG); huGAPDH-fw (ACTAGGCGCTCACTGTTCTC), -re (AATACGACCAAATCCGTTGACTC). Statistical analysis was performed using GraphPad Prism 8. For details refer to *Figure Legends*.

### Fluorescent immunolabeling and RNAscope

Human lung tissue and ALI culture membranes were fixed in 4% paraformaldehyde for 10 min at RT prior to paraffin embedding and cutting (tissue and vertical membrane sections). Sections were deparaffinized, blocked in 5% BSA/PBS/0.1% Tween 20 (one hour RT) and immunolabeled as previously described[4]. For RNAscope, slides were treated with protease plus (Advanced Cell Diagnostics) at 40° C for 30 min in a HybEZ hybridization oven (Advanced Cell Diagnostics) after antigen retrieval. Probes directed against hGPR87 (Advanced Cell Diagnostics, # 471861) mRNA and control probes were applied at 40°C in the following order: target probes, preamplifier, amplifier, and label probe for 10 minutes, and immunolabeled for basal cell markers (see above). Horizontal membrane immunolabeling (fixed in 4% paraformaldehyde, not paraffin embedded) was performed as previously described[10]. Nuclei and cell integrity were visualized by phalloidin and DAPI staining. Membranes were mounted on glass slides using Fluorescent Mounting Medium (Dako). Images of lung tissue sections were acquired using a Zeiss Axio Imager M2 (processed by Zen 3.3) or an Olympus VS-120 whole slide scanner using a 40x lens for RNAscope images. ALI cultures images were acquired using a Leica TCS SP8 confocal microscope and processed by ImageJ (NIH).

### Antibodies

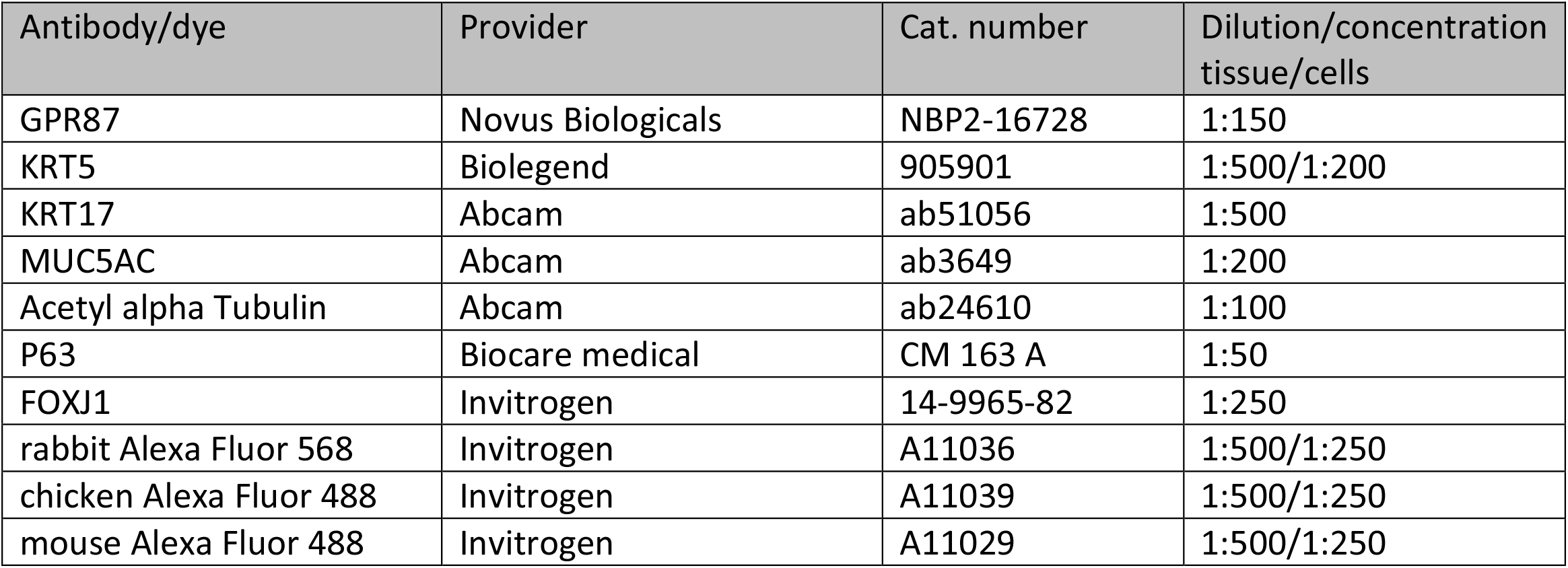
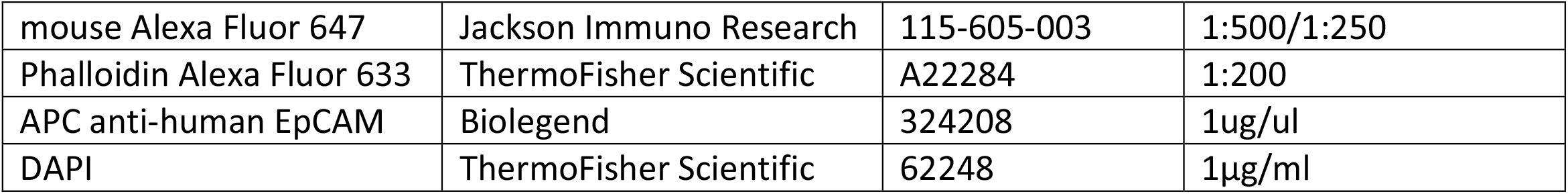

## Results

To identify potential novel epithelial cell populations in the IPF lung, we generated single cell transcriptomes from EpCAM^+^ cells isolated from parenchymal lung tissue from 3 IPF patients and 3 age-matched healthy donors. Using the 10x Genomics platform, we generated a dataset of 47,881 cells and found 10 distinct cell clusters, including main progenitor cell types of the alveolar region and distal airways as well as rare cell types, such as suprabasal cells, recently reported in the healthy lung[11] (A). Cells from both conditions were found in all clusters with differentially distributed clusters between healthy and IPF (B). In line with previous single cell data[6-8], ciliated cells were found mostly in IPF and ATII cells in non-diseased lungs, further underpinning a loss of ATII cells due to distal bronchiolization in IPF. Honeycomb cysts are an important histopathological criteria for the diagnosis of IPF, however, mechanistic insight in the process of bronchiolization and remodeling of the terminal bronchiole in IPF, remains largely unexplored. To shed light into cell populations potentially contributing to honeycomb cysts, we analyzed differentially expressed genes in all epithelial clusters and found cytokeratins such as *KRT6A, KRT5* and *KRT14* among the most upregulated genes in IPF (C). KRT5 is a well-characterized marker of basal cells and KRT5^+^ cells strongly accumulate in distal IPF lung tissues, mostly in areas of honeycombing[3, 4, 12]. We thus analyzed genes that correlate with KRT5 in IPF epithelial cells, and next to basal cell markers such as *KRT17* and known profibrotic genes such as *MMP1*, the highest correlation was observed with *GPR87*, a G-protein coupled receptor with unknown function in IPF and basal cells (D). *GPR87* was upregulated in several epithelial cell types in IPF, mainly in basal and suprabasal cells (E) in our data set, and confirmed with a published dataset[8] (F). We focused on GPR87 for several reasons: First, it belongs to the class of G-protein coupled receptors, which are intensively studied drug targets with attractive pharmacological accessibility. Second, although classified as an orphan receptor, profibrotic ligands have been discussed, such as lysophosphatidic acid[13]. Third, GPR87 has been linked to aberrant cell cycle control[14, 15], which is a feature of epithelial reprogramming in IPF[1]. Therefore, we aimed to investigate if GPR87 regulates epithelial cell reprogramming within distal IPF bronchioles.

We confirmed GPR87 epithelial expression and distrubution within the IPF lung *in situ* by RNAscope for *GPR87* RNA and immunolabeling with GPR87 antibody in distal IPF tissue sections. Notably, *GPR87* was detected in KRT5^+^ and KRT17^+^ basal cells of bronchiolar structures (G) and further enriched in clusters of KRT5^+^/p63^+^ basal cells in IPF lungs (H). GPR87 function was investigated in an air liquid Interface (ALI) cell culture model of HBECs, mimicking in vivo-like tissue with distinct cell types, including ciliated and secretory cells (I). GPR87 was expressed in KRT5^+^ basal cells of our human ALI culture (J). TGF-β treatment, inducing fibrotic epithelial reprogramming, led to increased *GPR87* expression in mature ALI cultures (K). This was consistent with the functional annotation enrichment analysis of our scRNAseq data, which revealed tissue development, keratinocyte differentiation, extracellular matrix remodeling, as well as TGF-β production, all indicative of altered epithelial airway differentiation and integrity, to be correlated with GPR87 (L). Morever, GPR87 overexpressing HBECs cultured at ALI displayed impaired differentiation evidenced by fewer ciliated cells and altered epithelial structure with larger apical surface area (M). Taken together, using scRNASeq of EpCAM enriched cells from IPF and donor lungs, we uncovered novel rare epithelial cell subpopulations in the IPF lung and for the first time report GPR87 as a novel marker and potentially druggable target of KRT5^+^ basal progenitor cells likely contributing to bronchiole remodeling and honeycomb cyst development in IPF.

## Figure Legends

**FIGURE 1.**
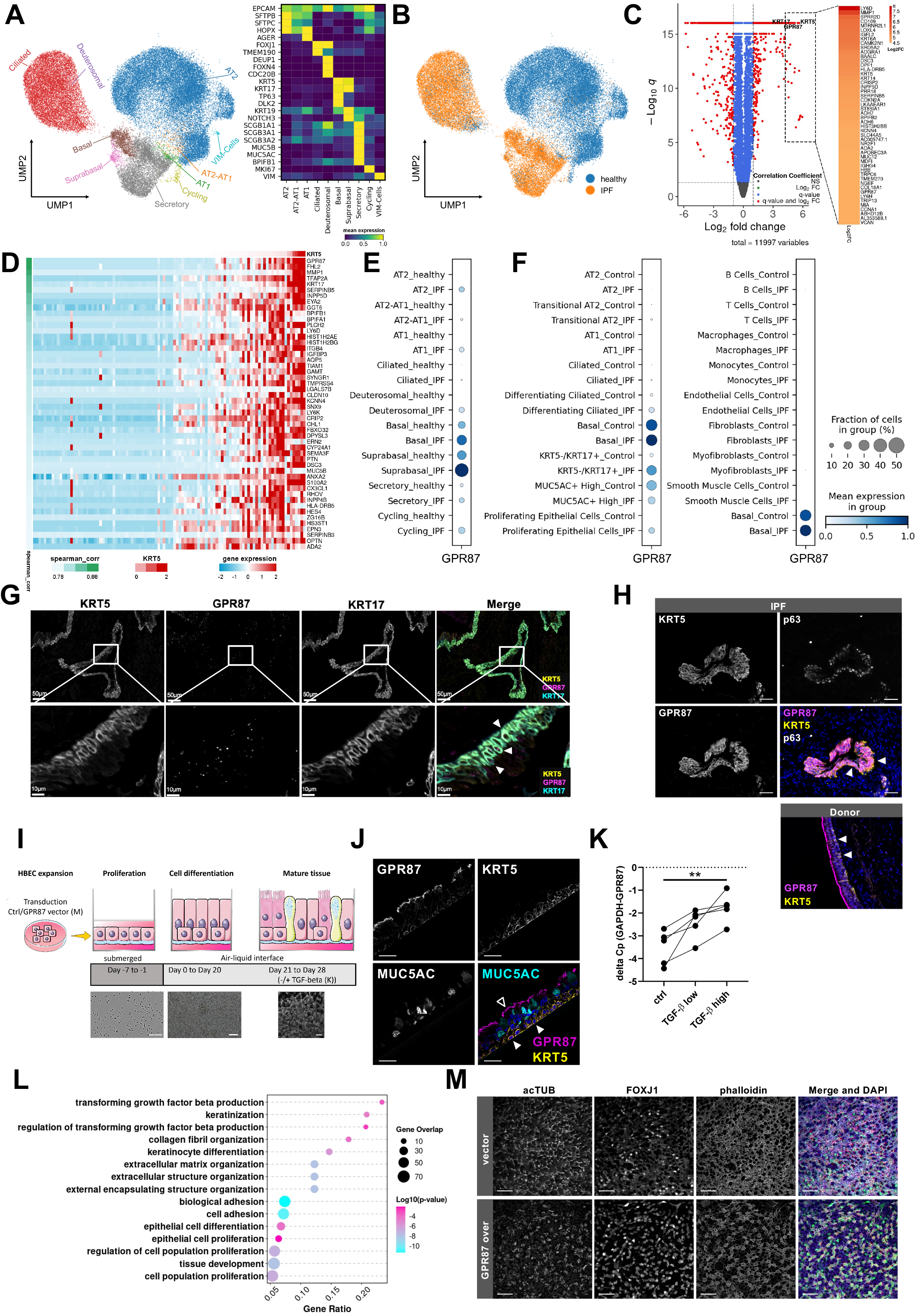
Accumulated basal progenitor cells highly expressing GPR87 localize within the IPF lung. **A)** Uniform manifold approximation and projection (UMAP) visualization shows unsupervised transcriptome clustering, revealing 10 distinct cell clusters. Heatmap shows highest expressed marker genes of each cluster. **B)** UMAP visualization showing distribution of healthy donor and IPF cells to different clusters. **C)** Volcano plot of differentially expressed genes (red, log2FC > 1, *p* < 0.05) in IPF EpCAM^+^ epithelial cells compared with donor samples, zooming in a gene set with top-50 fold change. **D)** Top 50 genes correlated with KRT5 are shown with gene expression level and Spearman’s rank correlation coefficient (spearman_corr). Dotplots show GPR87 expression in our **(E)** and a published dataset **(F)**, respectively. **G)** GPR87 mRNA was visualized by RNAscope, and combined with immunolabeling of basal(oid) cell markers KRT5 and KRT17. Higher and lower magnification of bronchioles are presented and indicated by respective bar sizes (10μm and 50μm). **H)** Lung tissue sections of IPF were co-immunolabeled for GPR87 and classic basal cell markers KRT5 and p63 (n=3), and healthy donor for GPR87 and KRT5 (n=2). Nuclei are visualized by DAPI staining. Representative double positive cells for respective markers are indicated by arrowheads. Bar size: 50µm; **I)** Scheme of HBEC isolation and ALI culture. HBECs were isolated from donor tissue and cultured on 6 cm dishes, coated with rat-tail collagen type I under submerged conditions. Cells were either transduced by virus for GPR87 overexpression (results (M)) and/or directly transferred to transwell plates and cultured on nylon membranes coated with collagen Type IV and with a pore size of 0.4 μm under submerged conditions for proliferation. Cells were airlifted (= day 0) and differentiated to a mature epithelium within 21 days. TGF-β treatment was performed at day 21 and every other day till day 28 (four times in total; results (K)). Shown are phase contrast images for dish cultured cells and early ALI (left, middle; bar sizes: 250μm, 100μm), and a confocal image of acTUB to visualize late ALI (mature epithelium, right, bar size: 25 μm). **J)** Vertical membrane sections of mature ALI cultured HBECs were immunolabeled for GPR87, basal cell marker KRT5 and secretory cell marker MUC5AC (n=2). Representative double positive cells for respective markers are indicated by arrowheads. Bar size: 25µm; (Cilia localization of GPR87, indicated by triangles, could be caused by non-specific antibody binding to cilia, however, based on our single cell dataset a positive GPR87 expression in other cell types might be considered but will be the subject of future investigations.) **K)** Airlifted donor HBECs were stimulated with low (2 ng/ml) and high (4 ng/ml) concentrations of TGF-β, as described in (I). GPR87 gene expression was assessed by qPCR in five independent donor cell lines. GAPDH was used as an housekeeper gene control. An one-way ANOVA and Dunnett’s multiple comparisons test was performed to determine statistical significance. ** p<0.05. **L)** Functional annotation enrichment analysis of GPR87 positive correlated genes reveals several categories of airway remodeling. **M)** HBECs were transduced with lentivirus containing the full ORF of GPR87 to generate a stable overexpression of GPR87 (GPR87-over). Empty backbone-vector alone was used as a control (vector). Cells were cultured on ALI till d21 and co-immunolabeled for acetylated tubulin (acTub) and FOXJ1. DAPI and phalloidin stainings were performed to visualize nuclei and cellular integrity. Representative images of an n=3 are shown. Bar size: 25µm.

## Notes

### Competing Interest Statement

The authors have declared no competing interest.

